# SoupX removes ambient RNA contamination from droplet based single-cell RNA sequencing data

**DOI:** 10.1101/303727

**Authors:** Matthew D Young, Sam Behjati

**Affiliations:** Wellcome Trust Sanger Institute, University of Cambridge; Cambridge University Hospitals NHS Foundation Trust, University of Cambridge; Department of Paediatrics, University of Cambridge

**Keywords:** scRNA-seq, Decontamination, Pre-processing

## Abstract

**Background:** Droplet based single-cell RNA sequence analyses assume all acquired RNAs are endogenous to cells. However, any cell free RNAs contained within the input solution are also captured by these assays. This sequencing of cell free RNA constitutes a background contamination that confounds the biological interpretation of single-cell transcriptomic data.

**Results:** We demonstrate that contamination from this ‘soup’ of cell free RNAs is ubiquitous, with experiment-specific variations in composition and magnitude. We present a method, SoupX, for quantifying the extent of the contamination and estimating ‘background corrected’ cell expression profiles that seamlessly integrate with existing downstream analysis tools. Applying this method to several datasets using multiple droplet sequencing technologies, we demonstrate that its application improves biological interpretation of otherwise misleading data, as well as improving quality control metrics.

**Conclusions:** We present ‘SoupX’, a tool for removing ambient RNA contamination from droplet based single cell RNA sequencing experiments. This tool has broad applicability and its application can improve the biological utility of existing and future data sets.

**Key Points:**

- The signal from droplet based single cell RNA sequencing is ubiquitously contaminated by capture of ambient mRNA.
- SoupX is a method to quantify the abundance of these ambient mRNAs and remove them.
- Correcting for ambient mRNA contamination improves biological interpretation.

## Introduction

Droplet based single-cell RNA sequencing (scRNA-seq) has enabled quantification of the transcriptomes of hundreds of thousands of cells in single experiments [1, 2]. This technology underpins recent advances in understanding normal and pathological cell behaviour [3, 4, 5, 6, 7, 8]. Moreover, large scale efforts to create a ‘Human Cell Atlas’ critically depend on the accuracy and cellular specificity of the transcriptional readout produced by droplet based scRNA-seq [9, 10].

A core assumption of droplet based scRNA-seq is that each droplet, within which molecular tagging and reverse transcription take place, contains mRNA from a single-cell. Violations of this assumption, which may distort the interpretation of scRNA-seq data, are common in practice. Clear examples include droplets that contain multiple cells (doublets), and empty droplets. Attempts to detect and remove doublets are an active area of research [11, 12, 13].

Another phenomenon that violates this assumption is the sequencing of cell free RNA from the input solution, admixed with a cell in its enclosing droplet. It is recognised that these contaminating non-endogenous RNAs are present even within data sets of the highest quality [2]. Nevertheless, no systematic effort has been made to quantify and compensate for their presence. The prevailing strategy for addressing cell free RNA molecules is to assume their contribution is negligible. The one exception to this is the case of mixed genotype single cell experiments where the genotype can be used to identify contamination and remove it [14].

Here, we show that this ‘soup’ of cell free mRNAs is ubiquitous and non-negligible in magnitude. Since the character and extent of ambient mRNA contamination varies by experiment, with increased contamination in necrotic or complex samples, ambient mRNAs may significantly confound the biological interpretation of scRNA-seq data. We present SoupX, a method for quantifying the extent of ambient mRNA contamination whilst purifying the true, cell specific signal from the observed mixture of cellular and exogenous mRNAs.

In this paper we begin by briefly describing the SoupX method. Following this we consider a range of datasets, summarised in Table S1. We first investigate two “species mixing” datasets run on the Chromium 10X [2] and DropSeq [15] platforms, which allow us to directly identify contaminating mRNAs and test our method’s accuracy. We then demonstrate how SoupX can be applied in practice using a dataset of peripheral blood mononuclear cells (PBMCs) [2]. We further explore the biological benefits of SoupX using a complex ‘kidney tumour’ dataset, which consists of 12 kidney tumour biopsies [16]. Finally, we conclude with some general remarks about ambient RNA contamination and the effects of failing to account for its presence.

### The SoupX method

Droplet based scRNA-seq methods produce counts of unique molecular identifiers (UMIs) for genes in thousands of cells. The aim of a scRNA-seq experiment is to infer the number of molecules present for each type of gene within each cell from this data. However, the observed counts arise from a mixture of mRNAs produced by the captured cell and those present due to background contamination. SoupX aims to remove the contribution of the cell free mRNA molecules from each cell and recover the true molecular abundance of each gene in each cell.

The algorithm consists of the following three steps (summarised in Figure 1):

i. Estimate the ambient mRNA expression profile from empty droplets.
ii. Measure or set the contamination fraction, the fraction of UMIs originating from the background, in each cell.
iii. Correct the expression of each cell using the ambient mRNA expression profile and estimated contamination.

SoupX produces a modified table of counts, which can be used in place of the original count matrix in any downstream analysis tool.

**Figure 1.**
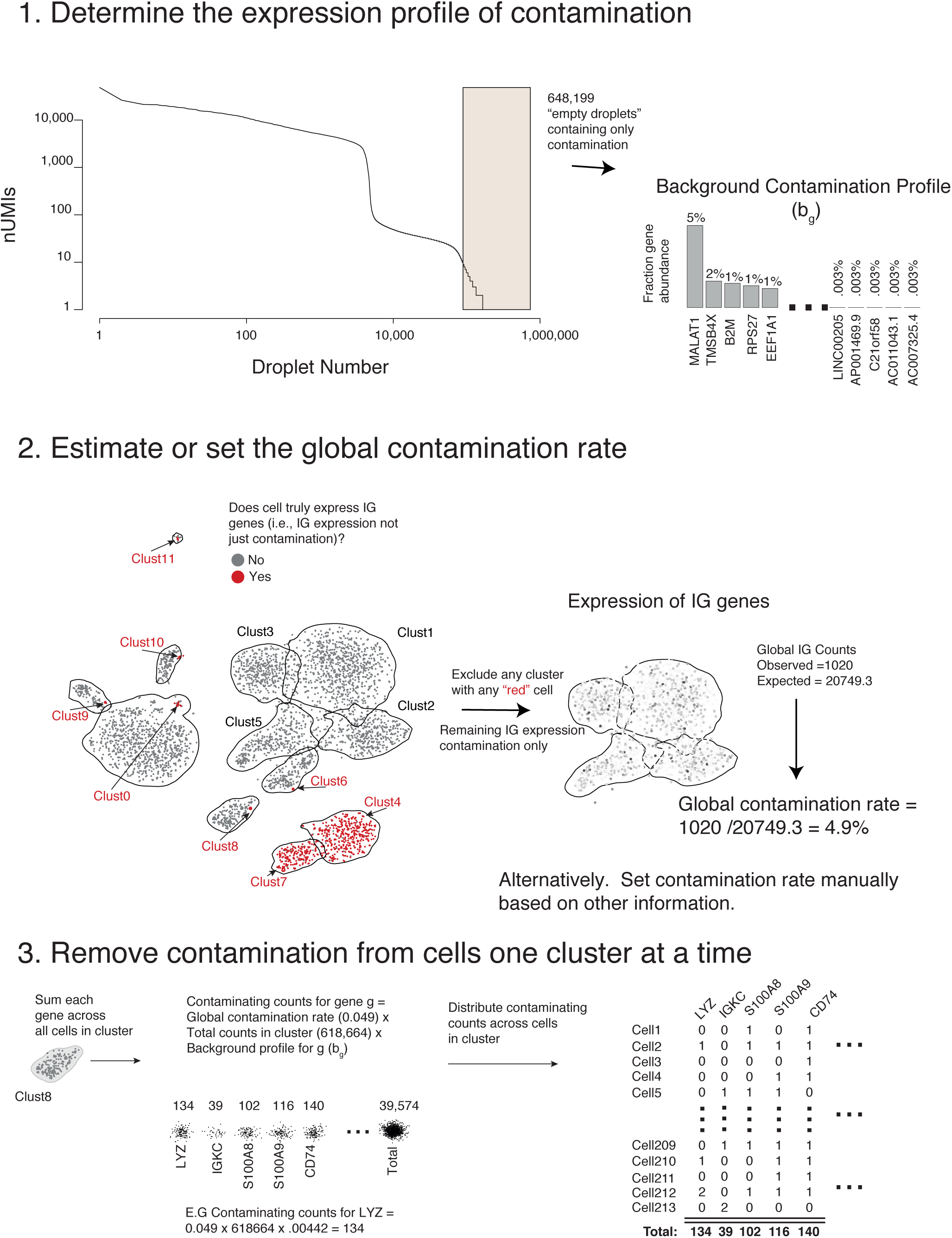
A visual summary of the SoupX method, using data from the PBMC dataset.

To estimate the background expression profile we consider all droplets with fewer than *N*_*emp*_ UMIs, which we assume unambiguously do not contain cells. The fraction of background expression from gene *g, b*_*g*_ is then given by,

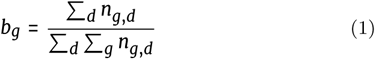

where *n*_*g,d*_ is the number of counts for gene *g* in droplet *d* and the sum over *d* is taken over all droplets with fewer than *N*_*emp*_ UMIs (Figure 1). The species mixing experiment allows us to compare how accurately *b*_*g*_ recapitulates the true background expression found within each cell, revealing that any value of *N*_*emp*_ < 100 produces a good correlation, with the best correlation given when *N*_*emp*_ < 10 (Figure S1).

The most challenging part of using SoupX is estimating or specifying the number of UMIs in each cell that are contributed by background contamination. In general, the observed number of UMIs for gene *g* in cell *c* is given by,

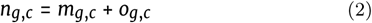

where *m*_*g,c*_ are the cell endogenous counts and *o*_*g,c*_ are the counts from the background. We assume the relative abundance of genes that make up the background does not differ between cells, which allows us to write,

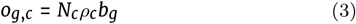

where *N*_*c*_ = ∑_*g*_ *n*_*g,c*_, and *ρ*_*c*_ is the background contamination fraction. In general *m*_*g,c*_ is unknown and what we are aiming to measure. However, there often exist certain combinations of genes and cells for which we can assume *m*_*g,c*_ = 0. It is the appropriate identification of these cells and genes that represents the greatest challenge in applying our method. How to do this in practice will be discussed below.

Given a set of genes/cells for which we can assume there is no cell endogenous expression (i.e., *m*_*g,c*_ = 0) we can calculate the cell specific contamination fraction,

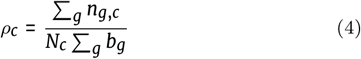

where the sum is taken across all genes in cell *c* for which it is assumed *m*_*g,c*_ = 0. Our method will optionally use clustering information to refine the set of cells for which it can be assumed that *m*_*g,c*_ = 0. If it can be shown for any cell *c* in cluster *P* that *m*_*g,c*_ > 0, then it is assumed that *m*_*g,c*_ > 0 for all *c* ∈ *P* (see Figure 1 and Section).

Having determined the contamination fraction *ρ*_*c*_ and the background expression profile *b*_*g*_, the cell endogenous counts are intuitively given by,

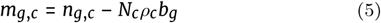

where *n*_*g,c*_ are the observed counts, *N*_*c*_ = ∑_*g*_ *n*_*g,c*_, and *b*_*g*_ and *ρ*_*c*_ are calculated as described above.

Although the intuition of Equation 5 is correct, in practice *m*_*g,c*_ is estimated by maximising a multinomial likelihood as described in Section. This procedure is further enhanced when cluster assignments are given, by performing the correction on counts aggregated at the cluster level, then distributing the corrected counts between cells in the cluster in proportion to their size (see Figure 1). This additional step helps overcome the sparsity of scRNA-seq data, which would otherwise make it impossible to distinguish a single count due to contamination from a single count due to endogenous expression in many circumstances.

The estimated value of *m*_*g,c*_ can then be used in place of *n*_*g,c*_ in any downstream analysis.

### Properties of ambient RNA

We next investigate the properties of ambient RNA contamination in data where ground truth is available, the “species mixing” experiments combining mouse and human cell lines [15, 2]. Figure 2A shows the relative abundance of human and mouse mRNAs in each droplet in the 10X data. Droplets containing human (top-right) and mouse (bottom-right) cells show that ∼ 1% of observed transcripts are cross species contamination. This rate of cross species contamination provides a lower bound on the total rate of ambient mRNA contamination as there will also be an additional contribution due to contaminating mRNAs from the same species (we later show the true contamination rate is ∼ 2%). A similar effect is seen in the Drop-Seq based species mixing data (Supplementary Figure S2). These observations demonstrate that cell free mRNA contamination is present even in highly controlled experiments.

**Figure 2.**
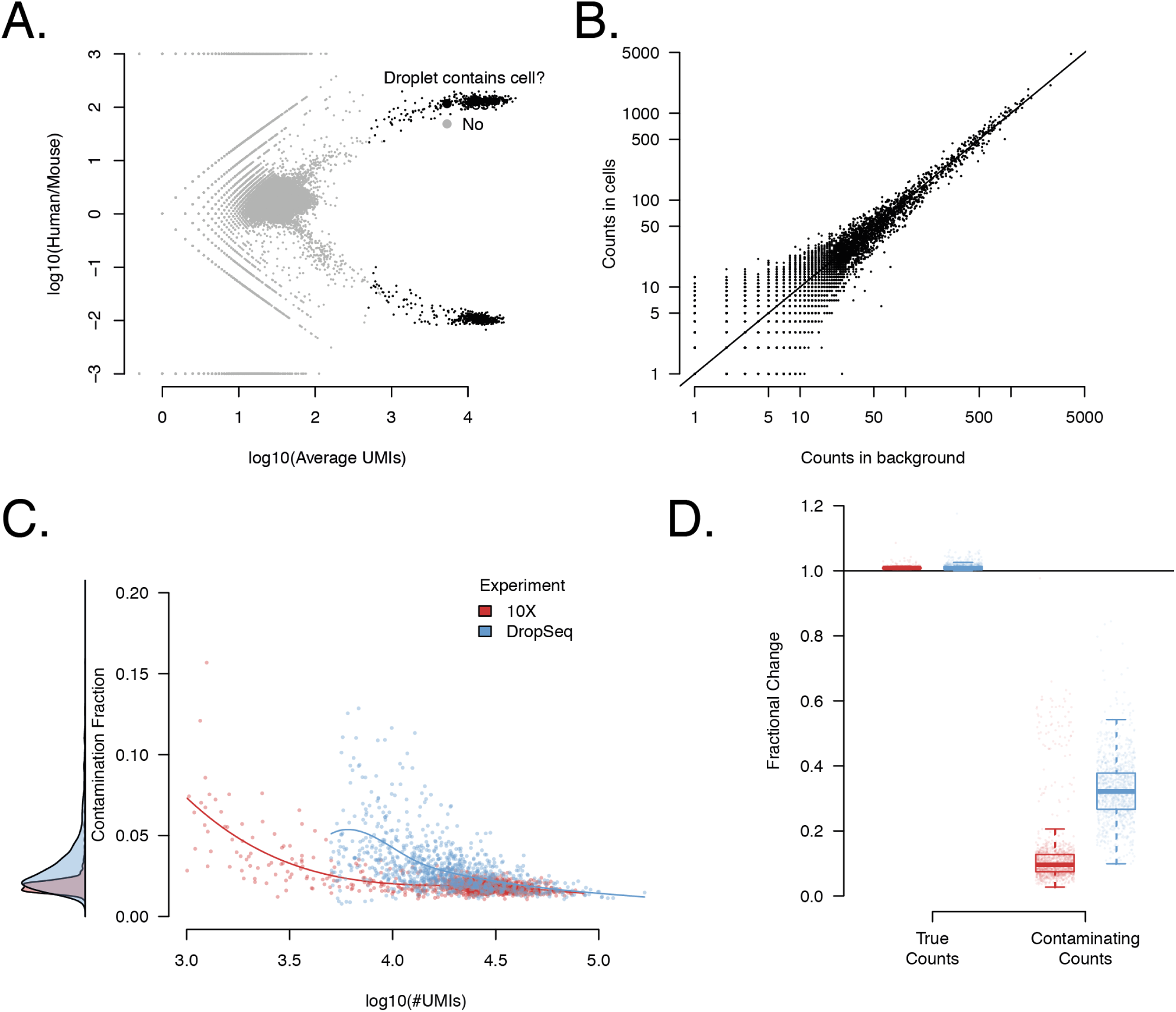
The properties of the cell free mRNA soup as determined using species mixing datasets. Panel A shows the log_10_ ratio of the number of UMIs mapping to human and mouse mRNAs for each droplet in the species mixing dataset (10X). Droplets determined to contain cells by cellranger are marked in black. Panel B shows the correlation of the counts in the background compared to counts averaged across cells for each gene. Counts have been sub-sampled so that the total number of counts in the background and averaged cell population is the same. Panel C shows the estimated contamination fraction as a function of number of UMIs in each droplet in individual cells in the species mixing dataset. Red/blue dots represent cells from the 10X/DropSeq experiments, respectively. The distribution on the left shows the marginal distribution across all cells. Panel D shows the fractional change in contaminating and genuine express levels after applying SoupX for the two technologies. The distribution across cells is summarised by boxplots, where the central line is the median, box boundaries are the 1st and 3rd quartiles and the whiskers extend to 1.5 times the interquartile range.

To investigate the composition of cell free mRNAs, we compared the aggregate expression profile of all droplets containing cells to all droplets with ≤ 10 UMIs, which we assumed to contain only ambient mRNAs. These two profiles were highly correlated in the 10X species experiment (Figure 2B) with a high correlation found in all other datasets considered (Pearson correlation 0.71 to 0.96, median 0.86; Table S2). The strength of the correlation implies that cell free contamination represents an approximately uniform sampling of the cells in the sequencing batch (i.e., channel).

Next we estimated the contamination fraction, the fraction of expression derived from the cell free mRNA background in each cell. In each cell we identify a set of genes that must have originated from the ambient mRNA; human transcripts in mouse cells and visa-versa. For these genes/cells it is assumed that *m*_*g,c*_ = 0 and the contamination fraction is calculated using Equation 4. Figure 2C reveals that there is little variation in the contamination fraction within a channel, in both the 10X and DropSeq data.

In most experiments there is less power to determine cell-specific contamination fractions and so SoupX assumes a constant contamination fraction within a channel. When clustering information is provided, the redistribution of counts from cluster level to individual cells, automatically removes more counts from contaminated cells, even when only a global estimate of the contamination is given (Figure S3; Section). Where a cell specific expression estimate is needed, SoupX employs a hierarchical bayes method to share information between cells (Section).

It may be hypothesised that the absolute number of contaminating mRNA molecules is the quantity that is approximately constant and that the contamination should vary with the number of mRNA molecules contributed by the captured cell. That is, that contamination fraction should vary as a function of cellular mRNA contribution, with the number of detected UMIs being a proxy for this. Consistent with this, Figure 2C shows that the greatest contamination occurs in droplets with the fewest UMIs. However, the contamination fraction is still approximately constant across most of the UMI range. This is likely a consequence of the fact that the capture efficiency of molecules in droplet based experiments varies by as much as an order of magnitude [17]. Thus variation due capture efficiency is likely to swamp variation due to “cell size” in most experiments, making constant contamination fraction a reasonable approximation.

To test the accuracy of SoupX in removing contaminating counts while retaining those due to endogenous expression we compared the fraction of expression from cross-species and within-species genes before and after SoupX contamination correction. This analysis revealed (Figure 2D) that mouse expression in human cells (and visa-versa) was decreased by at least a factor of two and usually an order of magnitude by the SoupX contamination removal in both 10X and DropSeq experiments. By contrast the fraction of expression derived from genes corresponding to the correct species was effectively unchanged for all cells.

## Application of SoupX to PBMC data

Next we tested our method on a data set consisting of PBMCs [2], measured in a single channel. We used the Seurat package [18, 19] to produce a tSNE representation of the data and annotated clusters of cells based on the expression of canonical marker genes (Figure 3A).

**Figure 3.**
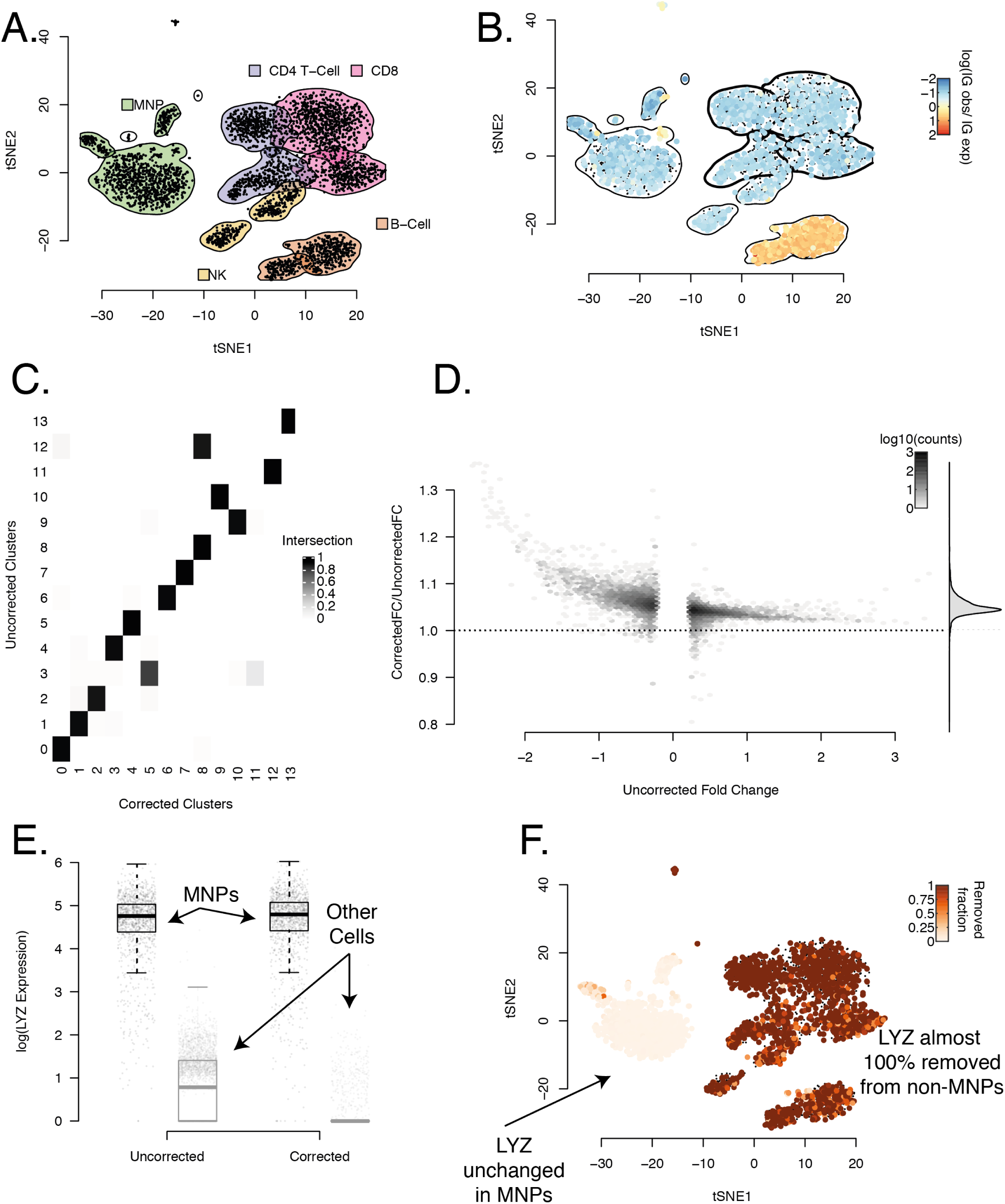
The PBMC dataset and how it changes when background correction is applied. Panel A shows a tSNE representation of the data, with cluster boundaries shown by density contours and shaded according to the cell type they represent. Panel B uses the same tSNE representation, but cells are now coloured by their rate of expression of IG genes compared to the rate at which IG is expressed in the background on a log_10_ scale. Positive values correspond to higher IG expression in a cell than in the background with values above significantly above 0 only possible if the cell endogenously expresses IG. The density contours of the clusters with no cell that endogenously expresses IG (as determined by a Poisson test) are marked in bold and used to estimate the global contamination ratio. Panel C shows the fraction of cells shared between clusters determined with the same parameters before and after application of SoupX. Panel D shows the improvement in marker specificity following application of SoupX. All genes that are markers of a cluster either before or after correction are identified and their expression log fold change relative to the clusters they do not mark is calculated before and after correction. The y-axis of this plot shows the fractional change in logFC for after applying SoupX for all genes. Genes are grouped into bins for ease of representation, with the number of genes in each bin given by the colour scale. The marginal distribution across all genes is shown on the right and the dotted line corresponds to no change in marker specificity after correction. Panel E illustrates the improvement in marker sensitivity for the gene LYZ, which is a marker for MNP cells. The corrected and uncorrected expression levels are shown split by cells labeled as MNPs and all others. Panel F shows this same change in expression on the tSNE map, where the colour scale represents the fraction of LYZ expression that has been removed by SoupX.

The main challenge in applying our method to this dataset was the selection of an appropriate set of genes that could be assumed to be unexpressed in some cells. That is, identifying the genes and cells for which it is safe to assume that *m*_*g,c*_ = 0.

To aid appropriate selection of such a gene set, we reasoned that the ideal genes for estimating the contamination rate would be ubiquitously present at a low level in all droplets due to high expression in the ambient RNA. They would also be present at a high level in those droplets that contain a cell endogenously expressing the gene, allowing us to unambiguously separate droplets with endogenously expressing cells (i.e., where *m*_*g,c*_ > 0) from those where the expression is solely due to contamination (*m*_*g,c*_ = 0).

Based on this reasoning, we developed a heuristic that ranks the 500 genes with the highest expression in the background by their bimodality of expression across all droplets in a channel. A plot based on applying this heuristic to the PBMC data shows the expression distribution across all cell containing droplets in the dataset (Figure S4). This heuristic suggests that immunoglobulin genes, such as IGKC and IGLC2, are both highly expressed in the soup and highly specific in their expression, making them good candidates for estimating the contamination fraction in this dataset. Note that this heuristic is intended only as an aid for selecting a biological sensible set of genes, not as an automated tool suitable for gene selection without manual intervention.

To select a precise set of cells for which we could use IG genes to estimate the contamination, we identified all cells whose IG expression was significantly greater than in the background contamination (Poisson test, FDR 0.05; Section). These represent cells endogenous expressing IG. We only used cells from clusters with no cells identified as endogenously expressing IG to estimate the contamination rate (Figure 3B). For the PBMC data, this identified IG expression in T cells as purely due to contamination and calculated a background contamination rate of ∼ 5%.

Having calculated the global contamination rate for the PBMC data, we then corrected the PBMC data for background mRNA contamination and reanalysed the data with Seurat using the same settings. Comparing cluster membership before and after correction revealed that the same number of clusters was identified, but some cells changed which cluster they belonged to (Figure 3C).

Next we identified marker genes for each cluster in both the corrected and uncorrected PBMC data using a Wilcoxon Rank Sum test and calculated the expression fold change between the cluster and all other cells. We compared the fold changes for the same genes in the same clusters before and after correction and found that correction for background contamination systematically increased the fold change contrast for marker genes (Figure 3D). That is, correction for background contamination made marker genes more specific to the cluster they were markers of. Furthermore, additional genes were found as markers in the corrected data, that were not identified in the uncorrected data.

As a specific example, we found that correction of ambient RNA contamination changes the pattern of expression of LYZ in the PBMC data (Figure 3E-F). This improved the specificity of LYZ as a marker gene for mononuclear phagocytes (Figure 3E) by removing its expression from all other cell types, while leaving its expression in mononuclear phagocytes unchanged (Figure 3F).

### Ambient RNA confounds interpretation in complex experiments

As a further test of the biological utility of our method we considered an experiment combining 7 kidney tumours processed across 10 channels (Table S1). As with the PBMCs, we analysed corrected and uncorrected data using the Seurat package; Figure 4A shows a tSNE plot of the uncorrected data. Haemoglobin genes were used to estimate the contamination fraction in most channels (Figure S5). This choice of geneset for estimating contamination was motivated by the ubiquitous presence of red blood cells (with red cell lysis forming part of the tissue treatment protocol) in these samples, together with the knowledge that HB genes are highly specific to erythrob-lasts.

**Figure 4.**
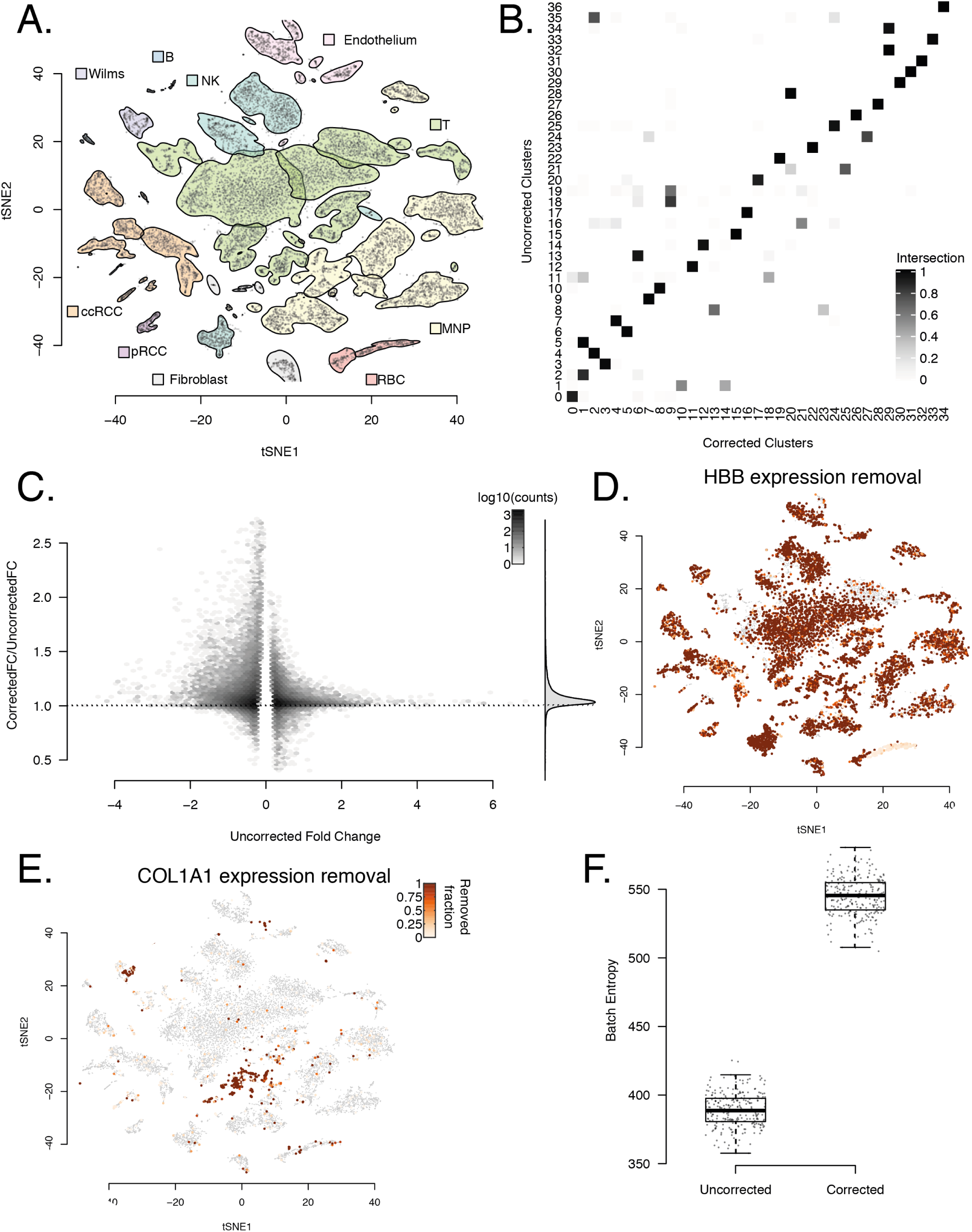
The application of SoupX to a complex, multi-channel dataset (the kidney tumour dataset). Panel A shows a tSNE representation of the data, with cluster boundaries shown by density contours and shaded according to the cell type they represent. Panel B shows the fraction of cells shared between clusters determined with the same parameters before and after application of SoupX. Panel C shows the improvement in marker specificity following application of SoupX. Note the different scale of the y-axis compared to Figure 3. All genes that are markers of a cluster either before or after correction are identified and their expression log fold change relative to the clusters they do not mark is calculated before and after correction. The y-axis of this plot shows the fractional change in logFC for after applying SoupX for all genes. Genes are grouped into bins for ease of representation, with the number of genes in each bin given by the colour scale. The marginal distribution across all genes is shown on the right and the dotted line corresponds to no change in marker specificity after correction. Panel D illustrates the improvement in marker sensitivity for the gene HBB, which is a marker for erythroblasts. The colour scale represents the fraction of HBB expression that has been removed by SoupX. Panel E the same as D but for COL1A1. Panel F shows the cross batch entropy before and after SoupX has been applied. The entropy measures the level of local mixing (100 nearest neighbours) for 100 cells selected from each cluster [20]

Applying SoupX and reanalysing the kidney tumour data revealed that in contrast to the PBMC data, many cells changed cluster and with the same clustering parameters two fewer clusters were identified in the corrected data (Figure 4B). Furthermore, we found that the expression ratio of marker genes between the cluster they mark and all other cells increased systematically after correcting for background contamination (Figure 4C).

We found that the correction of background contamination changed distribution of expression of many genes across cells in a way that would alter the biological interpretation. For example, while unlikely to be biologically misinterpreted, SoupX completely removes the expression of haemoglobin genes from all cells except erythroblasts (Figure 4D).

In other cases, the misattribution of gene expression to cell types that do not truly express them can lead to false conclusions. An example of this type is the cluster of T and mononuclear phagocytes (MNPs) in Figure 4A, E, which express the collagen genes COL1A1, COL1A2, and COL3A1 before background correction. The expression of collagen genes might be interpreted as evidence that the leukocytes are resident in the tissue. However, our method identifies that a high fraction of this expression is due to contamination (Figure 4E)

As the ambient mRNA expression profile is experiment specific, we reasoned that background contamination likely contributes a batch effects. That is, two identical cells captured in different experiments will appear different due to differences in their cell free RNA composition. We therefore calculated the cross-batch entropy of the Kidney tumour data before and after background correction [20] This analysis shows that the batch mixing entropy is increased after background correction, indicating better mixing between samples (Figure 4F).

## Discussion

We have shown that cell free RNA is omnipresent in droplet based scRNA-seq data and have proposed a method to identify, quantify and remove its contaminating effect. We find that accounting for contamination improves the specificity of marker genes, identifies new markers, and is essential for the correct biological interpretation of complex experiments.

We have shown potential misinterpretations of kidney tumour data driven by ambient mRNA contamination, but examples are sure to abound in other tissues. For instance, in endocrine tissues, it is crucial to understand which cell types secrete a particular hormone. The miassigned expression of even a single hormone gene can fundamentally change how investigators think about the cell type. Such problems will become increasingly common as efforts to compare similar cell types across tissues progress.

The main limitation of our method is the need to specify a set of genes for which it is safe to assume there is no cell endogenous expression. That is, the user must specify a set of genes and cells where it is safe to assume that the only source of expression for these genes is from background contamination. The expectation is that biological knowledge of the experiment being performed will guide this choice.

For example, solid tissue experiments are frequently highly contaminated with red blood cells and red cell lysis is used to prepare the samples [16]. As such, haemoglobin genes are often ubiquitously present in the background. Furthermore, erythroblasts are the only cells that produce haemoglobin under normal physiological conditions, so for the set of haemoglobin genes, it is safe to assume that there is no cell endogenous expression for cells that are not erythroblasts. Finally, erythroblasts express haemoglobin genes in such extreme abundance that they can be trivially identified by comparing the ratio of observed haemoglobin genes to that present in the background contamination (Figure S5)). These properties make haemoglobin genes a sensible choice for most solid tissue experiments.

In general the task of selecting a set of genes and cells will depend on the specifics of an experiment. We provide a number of heuristics that looks for bimodal expression amongst the genes highly expressed in the background contamination.

Having identified a useful set of genes, we provide a procedure for conservatively identifying which cells do not endogenously express the specified gene set. This procedure aggressively excludes clusters where there is clear evidence that at least one cell has expression of the target gene set exceeding what would be possible even in the extreme case of 100% contamination. This procedure can be made more or less conservative as needed either by changing the number of clusters considered, or by changing the cut-offs for the statistical test for extreme expression.

However, there will likely be cases where such genes cannot be identified. In such circumstances, SoupX may still be used by manually setting a global contamination rate, based on the experimenters best guess and contamination rates of similar experiments. For most applications the consequences of manually setting an unrealistically high contamination rate are likely to be minimal. Setting a higher global contamination rate than is realistic will decrease slightly the expression levels of genes that are truly specific to a cell, but as the relative abundance of cell specific genes are by their nature higher than in the background, true marker expression will never be completely removed.

SoupX has been designed to prevent the erroneous removal of genuine expression, however in many applications it may be preferable to over-correct for background contamination and remove a small amount of genuine signal in order to ensure that all the background contamination has been removed. Furthermore, we find that our method is robust to small inaccuracies in the estimation of the global contamination rate (Figure S3).

To make our method easily applicable, we provide an R package, SoupX, which can be used to estimate and remove this contamination. The output of this package is a corrected table of counts, which can be used as input for standard workflows. We envision background correction forming a standard part of droplet based scRNA-seq analysis pipelines.

## Competing interests

The authors declare that they have no competing interests.

## Author’s contributions

M.D.Y. conceived the project, developed the method and wrote the manuscript. S.B. contributed to the method development.

## Acknowledgements

We thank William Heaton and Valentine Svensson for discussions about droplet sequencing; Sarah Teichmann for discussions about the methodology; Sarah Teichmann, Aaron Lun and Manasa Ramakrishna for comments and review of the manuscript and Manasa Ramakrishna for improvements to the figures and their layout. Martin Prete for help with creating a docker version of the code. We thank Justin McManus and Mia Jaffe for discussions about all aspects of the paper, particularly around ways to automate estimation of contamination.

## Data availability

The 10X species mixing dataset was the mixture of the human cell line 293T and the mouse cell line 3T3 described in [2]. We used the data mapped and quantified using Cell Ranger 1.1.0 from https://support.10xgenomics.com/single-cell-gene-expression/datasets/1.1.0/293t_3t3. The dropseq species mixing data was obtained from [15], specifically SRR1748411. The PBMC data was taken from [2]. The ‘kidney tumour data set’ was taken from [16]. The mapped data sets supporting the results of this article are available in the GigaDB repository, [To be determined].

## Code availability

An implementation of the SoupX method described here can be found at https://github.com/constantAmateur/SoupX. The code used to generate the results presented in this manuscript can be found here https://github.com/constantAmateur/ambientRNA_paper. A docker container, containing all code and data needed to generate the results in this paper can be obtained from https://hub.docker.com/r/constantamateur/soupxpaper.

## Supplementary Materials

### Notation

We refer to the observed counts in a droplet *c* for gene *g* as *n*_*gc*_. Sums taken over a variable are represented with a., so *n*_.*c*_ = ∑_*g*_ *n*_*gc*_; the sum over all genes for droplet *c. m*_*gc*_ represents the cell endogenous counts present in a droplet *c*, for gene *g*. Similarly, *o*_*gc*_ represents the other counts in a droplet, contributed by background contamination.

We denote the fractional abundance of gene *g* in the soup or ambient mRNA background as *b*_*g*_. This is defined such that *b*_.·_= 1. *ρ*_*c*_ represents the background contamination fraction in droplet *c*, defined as *o*_.*c*_/*n*_.*c*_.

We define 𝒢 as the set of all genes that can be detected in a sequencing experiment.

### Choice of count distribution

The most appropriate model for count based data, such as the counts produced by scRNA-seq is a multinomial distribution. This distribution provides the probability of observing a given partition of *N* counts into *k* genes, given the relative probabilities of each gene *p*_*g*_.

In estimating how many counts to remove, where *N* tends to be small, we directly maximise the multinomial likelihood. However, in other cases we approximate the multinomial distribution as *k* Poisson distributions, which is an accurate approximation in the limit of large *N* and small *p*.

These distributions are commonly extended to include over-dispersion by using a Dirichlet multinomial or negative binomial distribution. Throughout this paper we ignore the effects of over-dispersion for computational expediency and because for most estimation procedures used the maximum likelihood estimator does not depend on the over-dispersion (e.g. estimating the mean).

### Detailed description of the SoupX method

As discussed in the manuscript, SoupX aims to remove the contribution of the cell free mRNA molecules from each cell. The algorithm consists of the following three steps:

i. Estimate the ambient mRNA expression profile from empty droplets.
ii. Measure the contamination fraction, the fraction of UMIs originating from the background, in each cell.
iii. Correct the expression of each cell using the ambient mRNA expression profile and estimated contamination.

SoupX produces a modified table of counts, which can be used in place of the original count matrix in any downstream analysis tool. This supplement provides the details for each of these three parts of the SoupX method.

#### Background expression profiles

To calculate the expression profile of cell free mRNAs, we assume that droplets with a very low UMI count contain only cell free mRNAs. As the number of droplets with low UMI counts is very large compared to the number of cells (∼ 10^6^ droplets versus ∼ 10^4^ cells), there is typically abundant power to accurately calculate the expression profile of the cell free background. Let 𝒟 denote the set of all droplets with a UMI count *α*_*l*_ ≤ *n*_.*c*_ ≤ *α*_*u*_. The background expression fraction for gene *g, b*_*g*_, is estimated as

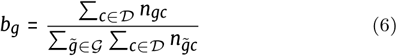

That is, *b*_*g*_ is the fraction of counts derived from gene *g* in the set of empty droplets 𝒟, normalised so that *b*_.·_= 1. In the species mixing data, we could directly measure the background contamination in droplets *with* cells. We used this gold standard to measure the accuracy of the estimated background as a function of the number of UMIs in the droplets used to estimate it (Figure S1). Based on this, we set *α*_*l*_ = 2 and *α*_*u*_ = 10 in this paper. We ignore droplets with 1 UMI to prevent errors in the droplet barcodes from contaminating our estimate of the background (although we find no evidence this is a problem for chromium 10X data). Different cut-offs may be more appropriate for different technologies, but we find good correlation between the expression profile of all droplets with less than ∼ 100 counts.

#### Calculating the contamination fraction

SoupX needs an estimate of the global contamination fraction present in a channel. This is generally not known in advance and must be estimated from the data or provided by the user. Our method approaches this problem by trying to identify a set of gene/cell pairs for which the cell endogenous expression can be assumed to be zero. That is, the task is to identify

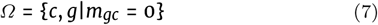

For genes and cells in *Ω* we see from equation 2 that *n*_*gc*_ = *o*_*gc*_. That is, the observed counts are purely due to background contamination.

In certain circumstances (e.g., a well annotated dataset with very specific marker genes) the set *Ω* may be able to be specified directly by the user. Where this is not the case, we construct *Ω* in a two-step procedure.

Firstly, a set of genes that are known to be very specific to a particular cell type and highly expressed in that cell type are identified. Typical examples of this are erythroblasts and haemoglobin genes or immunoglobulin genes and B cells. However, this set will depend on the experiment being performed, for example INS the insulin gene and pancreatic beta cells will work well in the pancreas and be useless elsewhere.

In cases where the choice of gene set is not obvious, we provide a heuristic plot that attempts to find genes highly expressed in the background with a bimodal expression profile across all cells. We focus on genes highly expressed in the background as in order to be useful, *Ω* must also have the property that ∑_*g,c* ∈*Ω*_ *n*_*gc*_ > 0. Put another way, the genes in *Ω* must be expressed at a reasonable level in the background so that we have sufficient counts to estimate the contamination fraction. Our heuristic therefore identifies the most useful genes as being highly expressed in the background, with a bimodal expression distribution across cells. This is motivated by the observation that the most useful genes are highly expressed in the cells they are specific to and have a low level of (contamination driven) expression elsewhere. This heuristic is intended only as a way to assist the development of a biologically motivated selection of an appropriate gene set. It is not suitable as a means for automating the SoupX method. Where automation is essential, it is better to manually specify a high global contamination (e.g. *ρ* = 0.15).

Having identified a set of genes suitable for estimating the contamination fraction, we next identify which cells definitively do not express these genes. Again, the ideal way to do this is to have a well annotated data set where this decision can be specified in a biologically motivated way. For example, when using haemoglobin genes to estimate the contamination fraction, cells annotated as erythroblasts can be excluded from *Ω* and other cell types included.

Where this is not possible we proceed by identifying all cells for which cell endogenous expression must be non-zero. These are the cells which are significant using a Poisson p-value under the null hypothesis that the *n*_.*c*_ counts are distributed in the same way as in the background distribution. That is, we identify all cells where *p*_*c*_ < 0.05 and

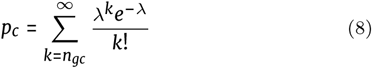

where *λ* = *ρ*_*max*_*b*_*g*_*n*_.*c*_ and *ρ*_*max*_ is the largest plausible contamination fraction (which is set to 1.0 by default). Put another way, this identifies all cells for which the fraction of counts derived from the genes of interest in the cell exceeds the fraction of counts for those genes in the background contamination.

We then cluster the data and exclude any cluster that contains a cell for which *p*_*c*_ < 0.05. This conservative approach helps ensure that the cells used to estimate the contamination are those cells with zero endogenous expression of the target genes. This approach can be made more or less conservative by adjusting the p-value threshold or clustering more or less finely.

Having constructed *Ω*, the set of gene/cell pairs with which to calculate the contamination fraction, we calculate the global contamination fraction for an experiment as

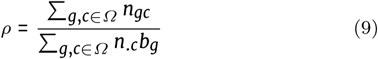

In most cases a global estimate of *ρ* is sufficient and we find little evidence of large cell to cell variability in *ρ*. Furthermore, in most cases the counts available to estimate the contamination within each cell, ∑_*g* ∈*Ω*|*c*_ *n*_*gc*_, is too low to provide an accurate cell level estimate.

For cases where there is a need to estimate cell specific contamination, we share information between cells using a Heirarcichal bayes model. Under the model:

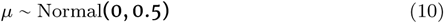

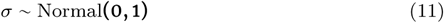

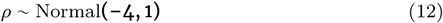

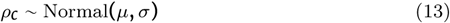

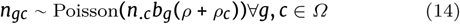

That is, the data is assumed to follow a Poisson distribution, with mean given by the expected background counts times a cell-specific contamination fraction. The cell specific contamination fraction is modelled as a global contamination, plus some perturbation whose prior distribution is normally distributed, the parameters of which are determined from the data.

#### Correcting cell expression profiles

Having calculated the expression profile for the background *b*_*g*_ and the contamination fraction *ρ*, we use this information to modify the table of counts and remove contaminating mRNAs, *m*_*gc*_. The obvious way to do this is by simply subtracting the contribution due to soup and setting *m*_*gc*_ = *n*_*gc*_ – *ρn*_.*c*_*b*_*g*_ (or *m*_*gc*_ = 0 if *n*_*gc*_ < *ρn*_.*c*_*b*_*g*_). Indeed, this is the maximum likelihood estimator of *m*_*gc*_ for Poisson distributed counts with mean given by equations 2 and 3.

However, following this approach will systematically under-correct the data as the only counts for which the data will be modified is those for which *n*_*gc*_ ≥ *ρn*_.*c*_*b*_*g*_. To correct for this, more than *ρn*_.*c*_*b*_*g*_ must be subtracted from those counts for which *n*_*gc*_ > *ρn*_.*c*_*b*_*g*_. The reason for this is that the data must be modelled by a distribution that takes into account the competitive nature of sequencing, such as the multinomial distribution. That is, we need a statistical model that will find not just the most likely amount of contamination in each gene separately, but will require that the total number of counts removed from each cell must equal *ρn*_.*c*_. The usual approach of modelling the counts for each gene/cell pair with a Poisson distribution approximates a multinomial distribution (similarly the often used negative-binomial distribution is an approximation of a Dirichlet-multinomial distribution). Therefore, the true problem we want to solve is to maximise the multinomial likelihood of *o*_*gc*_ (the contaminating counts for gene *g* in cell *c*) with multinamial *n* = *o*_.*c*_ = *ρn*_.*c*_, and probabilities given by *b*_*g*_, subject to the constraint that 0 ≤ *o*_*gc*_ ≤ *n*_*gc*_∀*g, c* (i.e., we cannot remove more counts than we observe).

We solve this problem by iteratively constructing *o*_*gc*_ by for each cell *c* as follows:

i. Initialise *o*_*gc*_ = min(*n*_*gc*_, *ρn*_.*c*_*b*_*g*_)∀*g* ∈ 𝒢
ii. Calculate the “unallocated counts” *F* = *n.c* – *o.c*
iii. Check if *F* < *ϵ*, if so stop.
iv. Set *o*_*gc*_ = min(*n*_*gc*_, *o*_*gc*_ + *ρFb*_*g*_)
v. Return to step ii.

where *ϵ* is a tolerance factor set to something small (such as 0.01).

This procedure is followed independently for each cell to produce modified counts. Where integer counts are required for downstream analysis, we round the corrected counts up to the nearest integer with probability given by the *m*_*gc*_ – ⌈*m*_*gc*_⌉.

#### Improving count removal using clustering

Where clustering of cells is provided, the above procedure can be improved by applying the correction procedure to counts aggregated within clusters. Doing this greatly increases the statistical power to distinguish between contamination and true expression. The value of *ρ* used for cluster *P* is calculated as

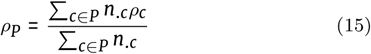

and the number of contaminating counts for each gene is calculated as above.

To redistribute the calculated contaminating counts to the single cell level, counts for gene *g* are distributed to each cell with weights given by *n*_.*c*_*ρ*_*c*_. This redistribution is done using the same iterative algorithm to ensure that a cell cannot be assigned more contamination for a gene than has actually been observed.

Cells that have a higher true contamination rate than the global average will have more non-zero counts in genes with high contamination than cells with a lower contamination rate than average. Because of this, the redistribution procedure described above will assign more contaminating counts to high contamination cells and fewer to low contaminating cells, even without this information being explicitly provided. This can be seen in Figure S3, where the effective cell level contamination rate implied by correcting at the cluster level and redistributing counts is highly correlated with the true cell level contamination rate.

### Processing of data sets

The DropSeq Species Mixing experiment was downloaded from the short read archive (SRR1748411) and quantified using Alevin [21] with a mixed human/mouse reference and the ‘forceCells’ flag set to 1 to include all barcodes.

For all 10X data sets, we used all droplets identified by cellranger as containing cells. In the species mix data, we removed any droplet with at least 1000 UMIs from *both* human and mouse genes, as these are likely doublets. For the DropSeq species mix experiment we set this threshold to 5000.

We used the Seurat package (http://satijalab.org/seurat/) to parse distinct cell types and marker genes from these preprocessed sequencing data. Raw counts of UMIs per gene in each cell were normalised using the Seurat::NormalizeData function, to implement the transformation

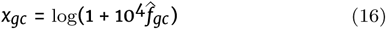

where 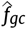 is the observed proportion of UMIs in droplet *c* from gene *g*; *x*_*gc*_ is the library-size normalised expression of gene *g* in droplet *c*.

Variable genes were identified using the Seurat::FindVariableGenes function with default parameters. Within Seurat, we subset the library-size normalized gene expression matrix on the variable genes, and we standardized the matrix so that the variable genes have mean 0 and standard deviation 1. We calculated the first 30 principle components of the standardized matrix, and the graph-based clustering algorithm implemented in Seurat::FindClusters evaluated the distance between cells in this 30-dimensional PCA volume. The t-SNE embedding was calculated using a perplexity of 30, and clusters were identified with the Seurat::FindClusters resolution parameter set to 1.

To identify genes specific to each cluster, we used the Seurat ‘FindMarkers’ function with default parameters.

These markers were then manually inspected and each cluster was assigned a cell type based on the comparison of these markers to the literature (particularly [22, 23, 24, 16]).

---

**Figure S1.**
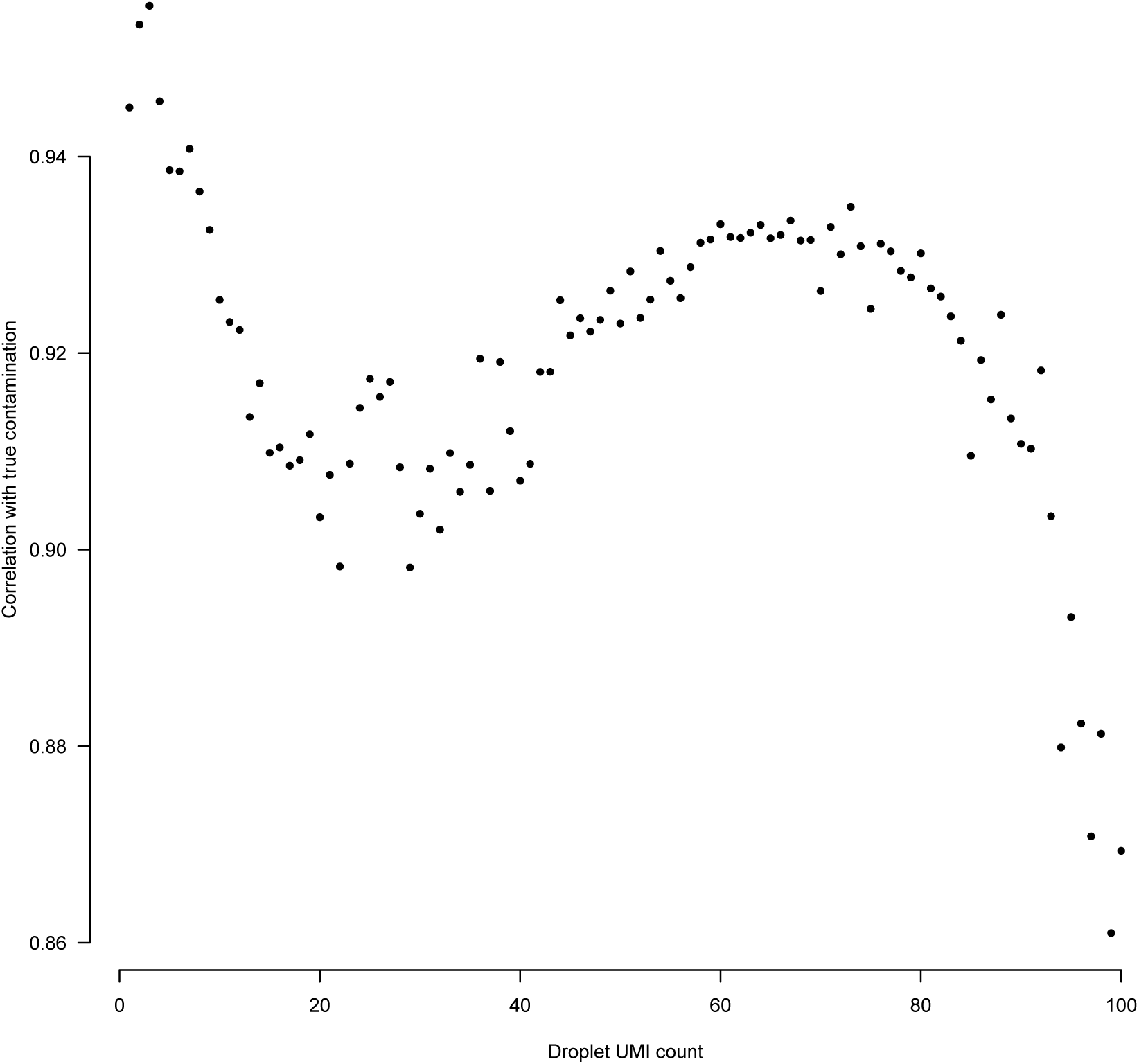
The correlation between “true background”, which is defined by aggregating across mouse transcripts in human cells and visa versa, with the background expression profile derived using only droplets with the total number of UMIs given on the x-axis.

**Table S1.** Sample information for the different datasets used in this paper.

| Dataset | Name | Platform | Num Channels | Sample type | Source |
| --- | --- | --- | --- | --- | --- |
| SpeciesMix | 10X | 10X | 1 | Human and Mouse cell lines | [2] |
| SpeciesMix | DropSeq | DropSeq | 1 | Human and Mouse cell lines | [15] |
| PBMC | PBMC | 10X | 1 | Peripheral blood mononuclear cells | [2] |
| KidneyTumour | Wilms1 | 10X | 6 | Wilms' tumour | [16] |
| KidneyTumour | Wilms2 | 10X | 3 | Wilms' tumour | [16] |
| KidneyTumour | Wilms3 | 10X | 3 | Wilms' tumour | [16] |
| KidneyTumour | PapRCC | 10X | 2 | Papillary cell renal cell carcinoma | [16] |
| KidneyTumour | RCC1 | 10X | 4 | Clear cell renal cell carcinoma | [16] |
| KidneyTumour | RCC2 | 10X | 4 | Clear cell renal cell carcinoma | [16] |
| KidneyTumour | VHL_RCC | 10X | 2 | Clear cell renal cell carcinoma | [16] |

**Figure S2.**
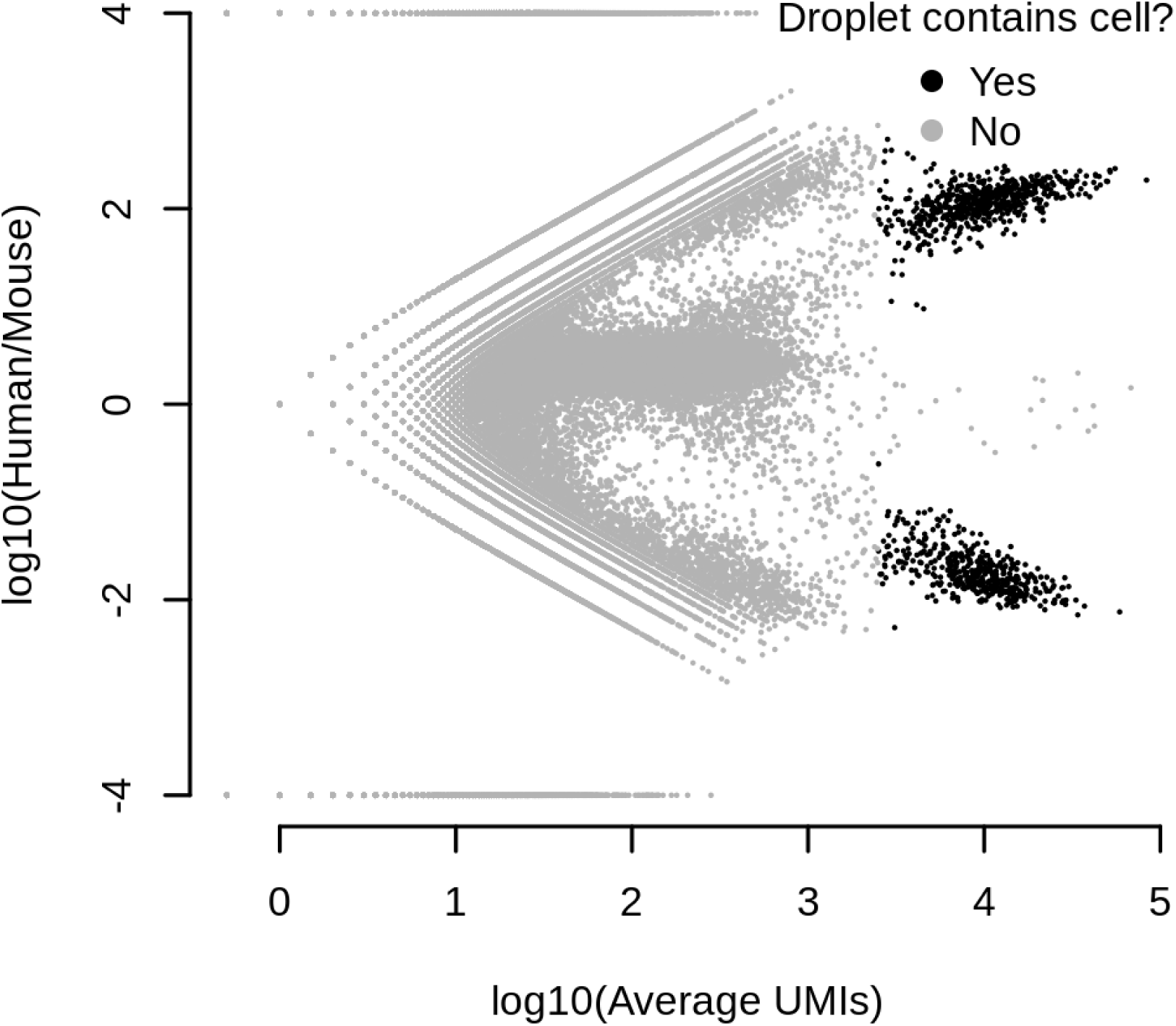
The ratio of human to mouse transcripts on a log_10_ scale (y-axis) for all droplets in the dropseq species mixing experiment. Droplets containing cells are marked in black. The x-axis gives the average number of UMIs between human and mouse for each cell.

**Figure S3.**
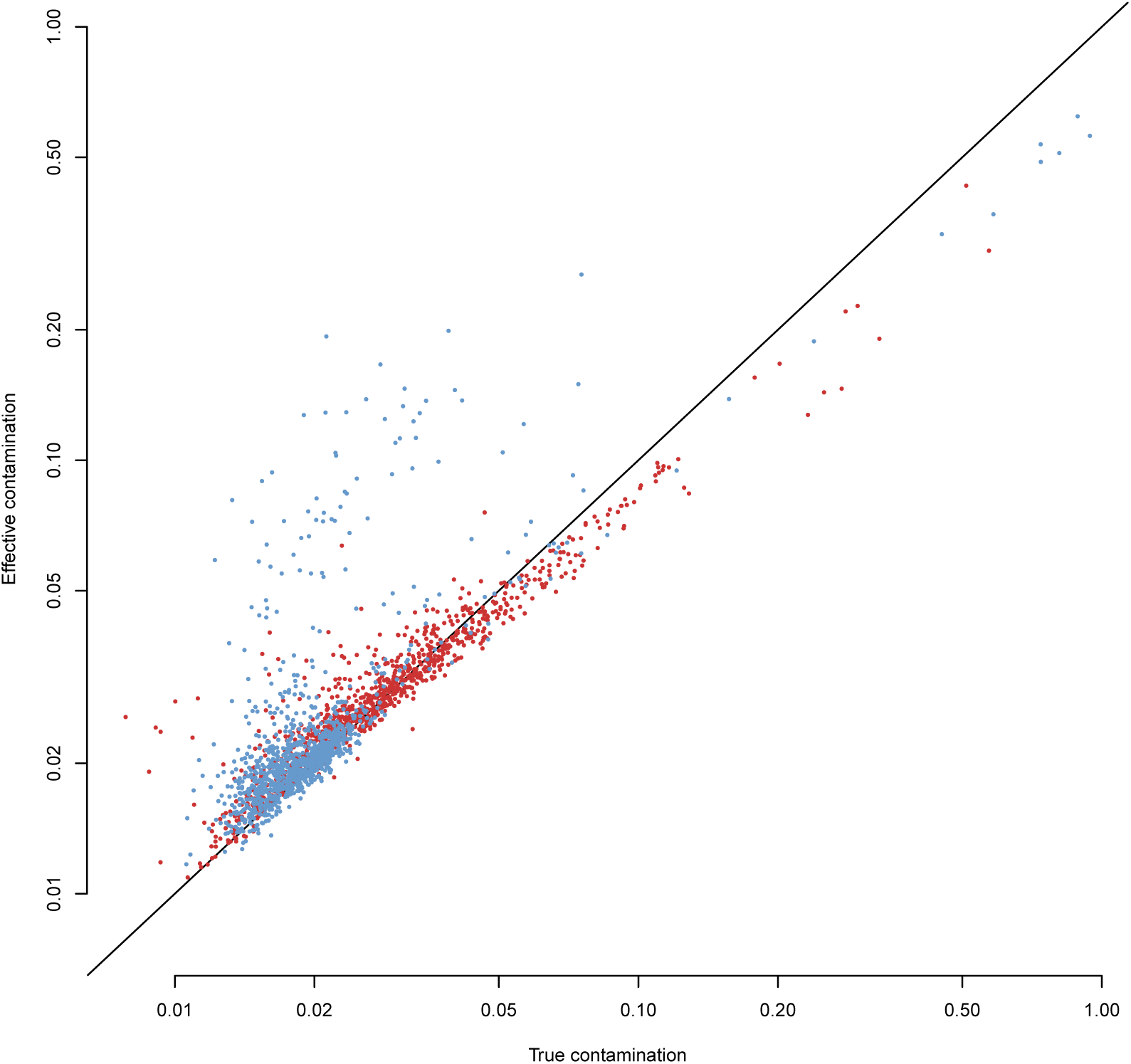
The x-axis gives the true contamination rate measured using the cross-species transcripts in each cell. The y-axis gives the effective contamination rate obtained by applying SoupX at the cluster level using a constant global contamination rate, calculated as the fraction of removed counts by the application of SoupX. The line shows perfect correlation and red/blue dots represent the 10X/DropSeq speciesmix experiments respectively.

**Figure S4.**
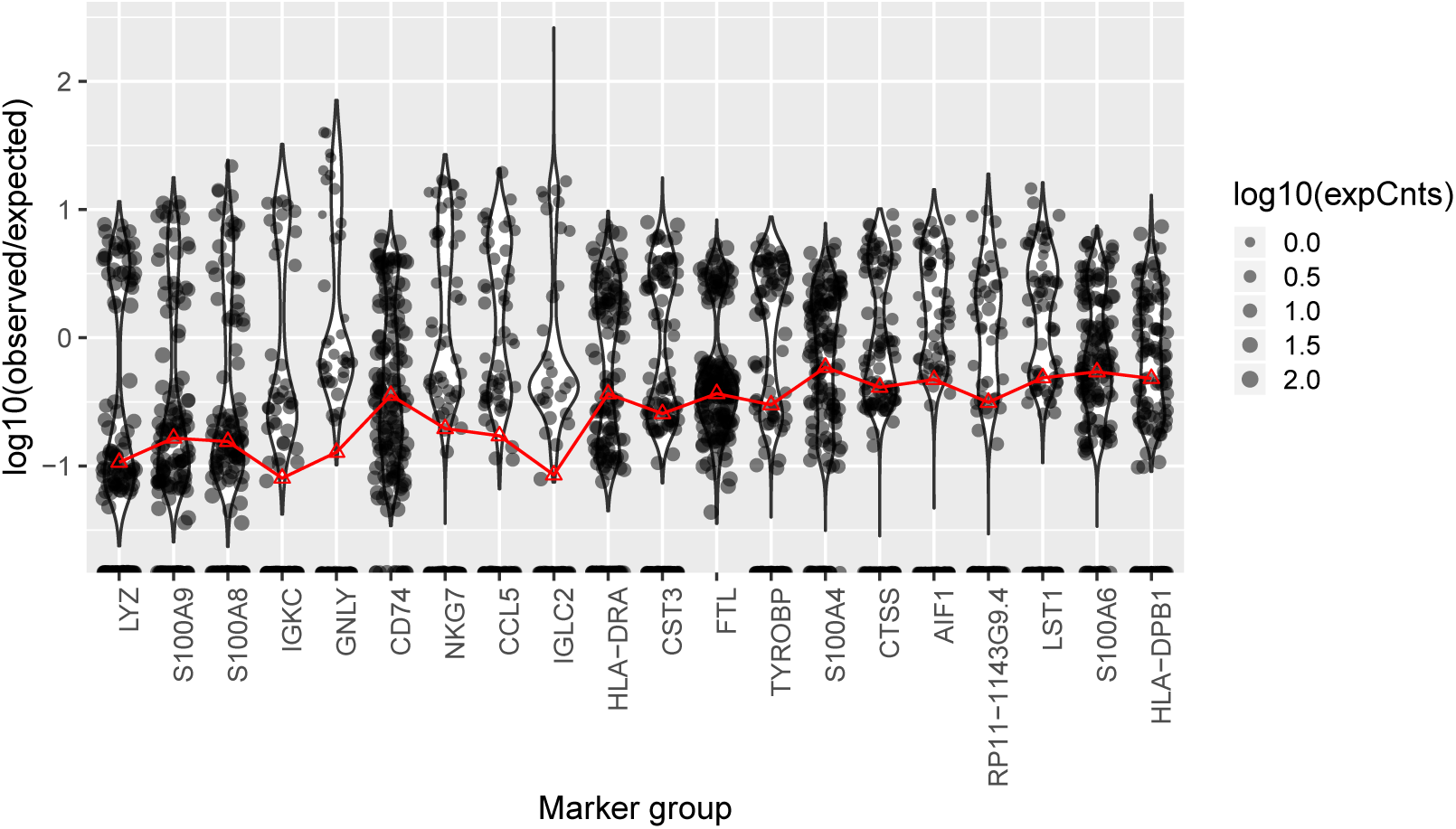
Distribution of expression relative to background for genes in the PBMC data. The red line indicates the global estimate of the contamination fraction that would be obtained if just that gene were used to estimate contamination. Genes which are most useful for contamination estimation have a bimodal distribution, with cells genuinely expressing the gene yielding a value on the y-axis greater than 0 and cells that do not express the gene having a value clustered around the true contamination rate.

**Figure S5.**
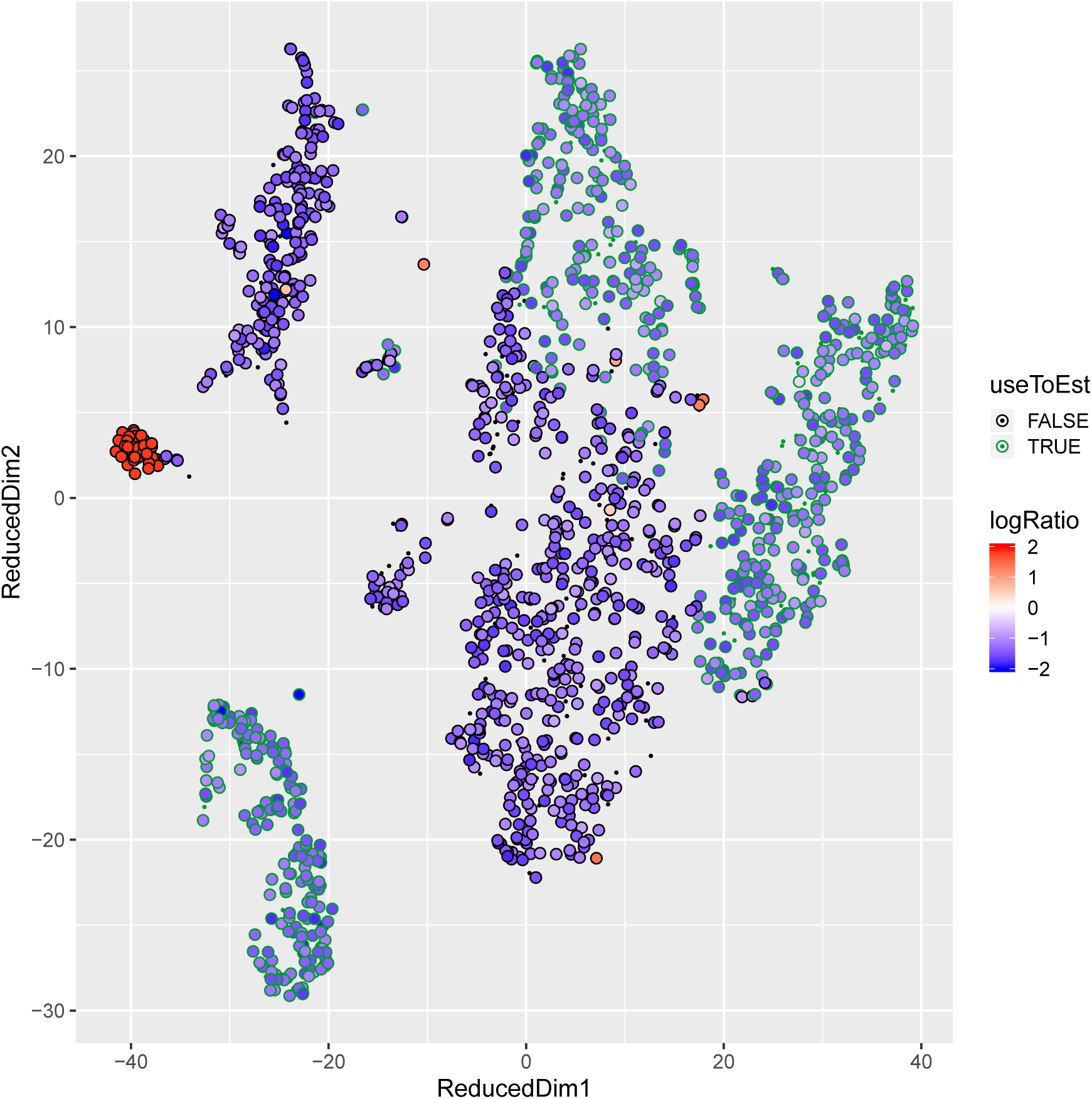
The fractional expression of haemoglobin genes in each cell, relative to the rate of expression in the background in one of the kidney tumour channels. This fraction is given by the colour of each point on a log scale. Points that have been determined to not endogenously express haemoglobin genes are marked with a green outline. The x and y axis are the tSNE coordinates supplied by cellranger for this channel.

**Table S2.** Pearson correlation coefficient between the background contamination profile and all cells in a channel averaged, after removing the genes above the 99th expression quantile.

| Dataset | Sample | Correlation |
| --- | --- | --- |
| SpeciesMix | 10X | 0.94 |
| SpeciesMix | DropSeq | 0.92 |
| PBMC | PBMC | 0.96 |
| KidneyTumour | Wilms3_Kid_T_ldc_1_1 | 0.92 |
| KidneyTumour | Wilms3_Kid_T_ldc_1_2 | 0.93 |
| KidneyTumour | Wilms3_Kid_T_ldc_1_3 | 0.93 |
| KidneyTumour | Wilms2_Kid_T_ldc_1_1 | 0.73 |
| KidneyTumour | Wilms2_Kid_T_ldc_1_2 | 0.73 |
| KidneyTumour | Wilms2_Kid_T_ldc_1_3 | 0.77 |
| KidneyTumour | Wilms1_Kid_R_ldc_1_1 | 0.75 |
| KidneyTumour | Wilms1_Kid_R_ldc_1_2 | 0.74 |
| KidneyTumour | Wilms1_Kid_R_ldc_1_3 | 0.85 |
| KidneyTumour | Wilms1_Kid_T_ldc_1_1 | 0.84 |
| KidneyTumour | Wilms1_Kid_T_ldc_1_2 | 0.86 |
| KidneyTumour | Wilms1_Kid_T_ldc_1_3 | 0.84 |
| KidneyTumour | VHL_Kid_T_ldc_1_1 | 0.82 |
| KidneyTumour | VHL_Kid_T_ldc_1_2 | 0.81 |
| KidneyTumour | RCC2_Kid_T_ldc_1_1 | 0.81 |
| KidneyTumour | RCC2_Kid_T_ldc_1_2 | 0.84 |
| KidneyTumour | RCC2_Kid_T_ldc_2_1 | 0.89 |
| KidneyTumour | RCC2_Kid_T_ldc_2_2 | 0.89 |
| KidneyTumour | RCC1_Kid_T_ldc_1_1 | 0.86 |
| KidneyTumour | RCC1_Kid_T_ldc_1_2 | 0.84 |
| KidneyTumour | RCC1_Kid_T_ldc_2_1 | 0.87 |
| KidneyTumour | RCC1_Kid_T_ldc_2_2 | 0.87 |
| KidneyTumour | pRCC_Kid_T_ldc_1_1 | 0.94 |
| KidneyTumour | pRCC_Kid_T_ldc_1_2 | 0.94 |

